# Epithelial Modelling Platform: A Tool for Model Discovery and Assembly with the Physiome Model Repository

**DOI:** 10.1101/631465

**Authors:** Dewan M. Sarwar, Yuda Munarko, David P. Nickerson

## Abstract

In this paper we present a web-based platform enabling scientists to construct a novel epithelial transport model to investigate their hypotheses, aided by building on existing models discovered in the Physiome Model Repository (PMR). We have comprehensively annotated a cohort of epithelial transport models deposited in the PMR as a seeding collection of building blocks that are freely available for reuse. On the platform, users are able to semantically display models for visualization, graphical editing, and model assembly. In addition, we leverage web services from the European Bioinformatics Institute (EBI) to help rank similar models based on the suggestions provided by the platform. In addition to potential use in biomedical and clinical research, novice modellers could use our platform as a learning tool. The source code and links to a live demonstration of the platform are available at https://github.com/dewancse/ epithelial-modelling-platform.

## 1 INTRODUCTION

The IUPS Physiome Project (Hunter and Borg, 2003) and the Virtual Physiological Human (VPH) (Hunter et al., 2010) have introduced tools and open source software for scientists to conduct biomedical research (Garny et al., 2010; Cooper et al., 2010). For example, biomedical engineers and clinicians often utilize computational models to investigate experimental or clinical hypotheses that are difficult or expensive to achieve experimentally using animal or human subjects. Modern tools and technologies help them to quickly and precisely test such hypotheses which may involve disparate biophysical symptoms and/or mechanisms such as clinical observations, diseases, and drug actions through physiology pathways (de Bono et al., 2016).

In this paper, we present a web-based platform, the Epithelial Modelling Platform (EMP), which will enable biomedical engineers and clinicians to discover epithelial transport models with a view to constructing a new epithelial model. This new model will help them to investigate specific research questions and hypotheses. To prepare a model discovery environment, mathematics-and physics-based epithelial transport models are integrated with biological information. Integration of such information in the context of biology is called model annotation. Model annotation captures semantic meanings of biological phenomena encapsulated in mathematics and thus hides underlying mathematical equations (Lister et al., 2009). In order to represent biological phenomena, model annotation makes use of a rich set of controlled vocabulary, i.e. ontologies (Courtot et al., 2011). This often requires multiple reference ontologies to represent such phenomena, which is called composite annotation (Gennari et al., 2011). Model annotation is a time-consuming process and thus requires automated process. For this, SemGen has been developed to annotate biosimulation models (Neal et al., 2018). We have deposited the annotated information and the models in the Physiome Model Repository (PMR) (Yu et al., 2011) where information is discoverable, accessible, and persistent. PMR follows FAIR (Wilkinson and et al., 2016) principles; i.e. information in PMR is Findable, Accessible, Interoperable, and Reusable (FAIR).

Prior research in (Galdzicki et al., 2011; Neal et al., 2013) demonstrates that model discovery can be performed by utilizing semantic querying technologies. Similarly, with the provision of modern technologies, knowledge discovery has introduced a range of tools and open source software such as DisGeNET (Piñero et al., 2015), ApiNATOMY (de Bono et al., 2014), JWS Online (Wimalaratne et al., 2004), Saint (Lister et al., 2009), semanticSBML (Krause et al., 2010), eSolve (de Boer et al., 2017), Chalkboard (Cook et al., 2007), Physiomaps (Cook et al., 2013). A recent initiative in (Cooper et al., 2015a), the Cardiac Electrophysiology Web Lab, has introduced an online simulation environment to compare simulation experiments. Such virtual experiments are analogous to *in silico* environment of *in vivo* and *in vitro* experiments, or wet lab experiments (Cooper et al., 2015b).

In addition to model discovery, we have assembled the discovered models on the platform with a view to constructing a new epithelial model. Model construction integrates components from various source models. SemGen can compose models in CellML (Cuellar et al., 2003) and SBML (Hucka et al., 2003) format. CellML and SBML are XML-based languages to describe biological processes. Specifically, SemGen internally converts models into OWL format and then does model composition (Hoehndorf et al., 2011). In contrast, we have composed models by utilizing libCellML (libCellML, 2019). libCellML is a library to serialize, validate, and instantiate a CellML model. A great deal of previous research on model composition is investigated in (Neal et al., 2015; Gennari et al., 2008; Neal et al., 2009; Krause et al., 2010; Coskun et al., 2013).

Here we present a model discovery and assembly approach which is a new contribution to PMR. In addition to potential use in biomedical and clinical research, novice modellers could use our platform as a learning tool. This platform would demonstrate how semantic web technologies and methodologies can contribute to educational purpose. In the section Materials and Methods, we present our current workflow of the EMP, the standards and software tools we have used to encode the computational models and semantic annotations for our exemplar cohort of models, and the technologies that enable the implementation of our platform. This platform, which makes use of the encoded models and semantic annotations to discover, explore, and visualize models, is discussed in the Results section. The discussion section summarizes the model discovery and assembly of the EMP with some concluding remarks.

## 2 MATERIALS AND METHODS

Fig. 1 presents an overview of the knowledge management platform being developed and utilized in this work. On the client side, content creation and editing applications like OpenCOR (Garny and Hunter, 2015) are used to encode computational models and associated information in computable formats. Such applications work on local workspaces, which can be synchronized with cloud-based repositories such as the PMR. Synchronization with PMR ensures persistent, resolvable, and discoverable identifiers for all resources in the workspace. For composite annotation, we used the SemGen tool and deposited this annotation as metadata in PMR (Sarwar et al., 2018). Web-based applications such as our EMP are then able to leverage the knowledge stored in PMR to discover models, and semantically assemble these models on the platform for visualization, graphical editing, and model assembly and construction.

**Figure 1.**
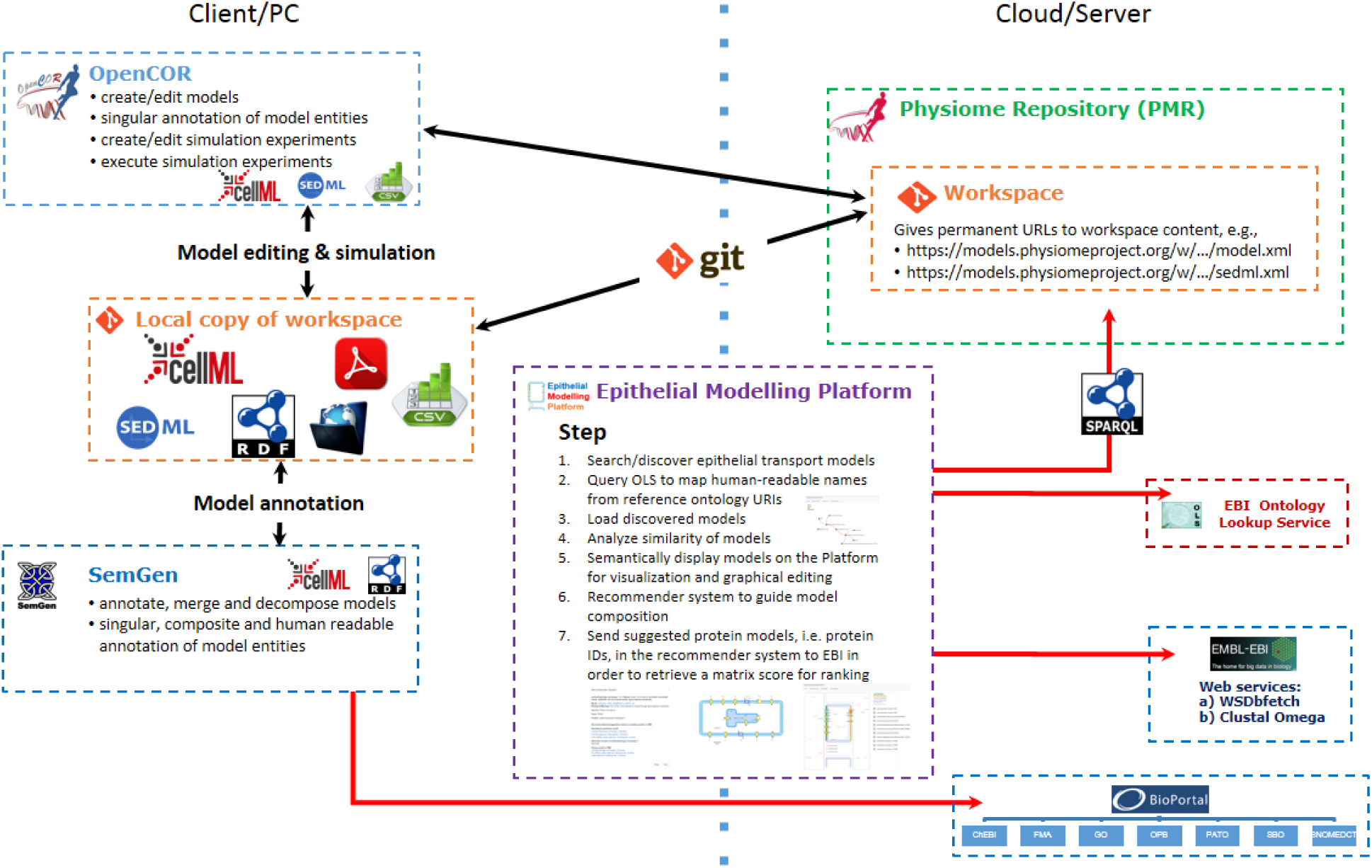
Illustrative example of current workflow for: 1) using OpenCOR to edit CellML models; 2) using SemGen to annotate the biological knowledge and deposit them in PMR; 3) using a git client to synchronize the local data to PMR; and 4) using our Epithelial Modelling Platform to discover CellML models and then semantically display these models on the platform for visualization, graphical editing, and model assembly and construction.

Our EMP has become a key demonstrator application of the knowledge management system. The aim was to discover models from the PMR and assemble these on the platform with the provision of the EBI ontology lookup service (OLS) (OLS, 2019) and web services as shown in Fig. 1. The seven key steps in this process are described as follows. (1) Search/discover models of interest using the information extracted from the annotations deposited in PMR. (2) In this case, the EBI OLS is used to map human-readable names to reference ontology identifiers. (3) The user will then select models from the discovered model set and will make a shortlist of models for visualization and graphical editing on the platform. Similar to (2), the EBI OLS is used for name resolution. (4) In the shortlisted space, we have provided an option for the user to find similar components between a set of selected models, enabling users to identify overlap between models or points of integration. (5) We then semantically visualize the shortlisted models on a scalable vector graphics (SVG), which is an XML-based two-dimensional interactive display format, display panel for graphical editing. Specifically, the shortlisted models are placed on a specific membrane or compartment based on the annotation in PMR. (6) A recommender system is implemented to help the user enhance and extend the model they are building by providing links, for example, to models of similar transporters or transporters of the same solutes in different membrane locations. Such user-editing will automatically change the default settings of the models on the platform. Suggestions in this recommender system are provided from the available annotated apical or basolateral membrane models. (7) The EBI web services are used to rank the available models in the recommender system.

Computational tools and standards have been evolved over the years by The COmputational Modeling in BIology NEtwork (COMBINE) (COMBINE, 2019) and the World Wide Web Consortium (W3C) in order to collaborate and disseminate tools and standards. Some of the tools and standards used in our EMP are described in (Sarwar et al., 2018).

## 3 RESULTS

We have developed the EMP to discover and explore computational models of interest, as well as to semantically visualize the discovered models on the platform for graphical editing and model assembly with a view to constructing a new epithelial model. In particular, this involves annotation of mathematical models with biological information and generation of metadata using the SemGen tool so that our EMP could discover models from the metadata. Annotation of a cohort of epithelial transport models have been discussed in (Sarwar et al., 2018), therefore, following section presents a detailed workflow of the EMP.

### Discover CellML Models

The first workflow of the EMP is to discover and explore models of interest extracted from the annotation in PMR. Fig. 2 shows a list of discovered models for a search query “flux of sodium” from PMR. From this list, the user can investigate various options such as CellML model entity – name of the model, component name and variable name; biological meaning deposited in PMR; protein name; and species and genes used during experiments of the associated models. In addition, the user can navigate to the “view” option to explore more information about a model. The user can make a shortlist of models by navigating through the “Add to Model” option and can return here in order to rediscover models.

**Figure 2.**
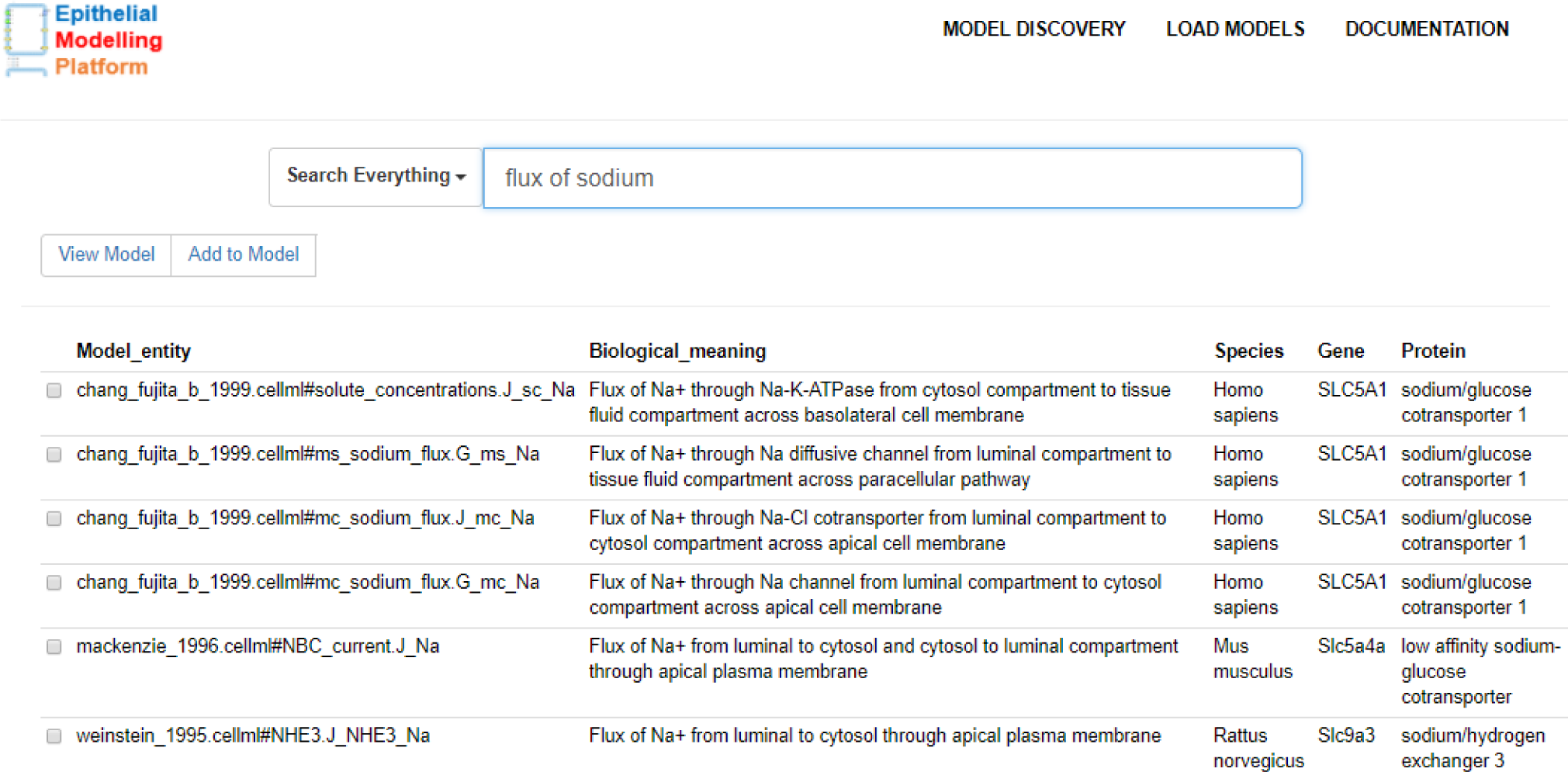
Model discovery interface to search for models in PMR that are relevant to the query “flux of sodium”. By querying the annotations stored in PMR, this interface retrieves components from the NHE3 (Weinstein, 1995), SGLT2 (Mackenzie et al., 1996), and an epithelial cell model (Chang and Fujita, 1999).

Users are able to enter human-readable search terms and phrases or choose to enter specific vocabulary terms to search for. When discovering resources in PMR relevant to a given search phrase, we first decompose the English prose into a series of semantic queries which are executed on the PMR knowledge base and ranked according to how well the discovered resource matches the initial search query. Continuing our example, the phrase shown in Figure 2 will best match any resource associated directly with a sodium flux, but other sodium-related resources will also be returned with a lower ranking in the search results. To achieve that, we have maintained a static dictionary with key and value pairs to map searched text to reference ontology terms. For example, for a search query “flux of sodium”, the model discovery interface will map flux and sodium to OPB and ChEBI ontology terms, respectively, as defined in the dictionary.

In addition to model discovery, users are able to add protein models by querying to the bioportal web service. This begins with entering a protein name in the “Add Model” interface of the EMP and then the bioportal web service returns a protein ID from the protein ontology (PR). Afterwards, the interface sends this protein ID to the OLS to retrieve species and gene names. Next, users enter the rate of concentration of solutes (i.e. flux) moving between compartments across apical or basolateral membrane. After filling in the required information, metadata will be generated and a CellML model will be serialized. In the future, we will implement a notification system in order to notify this new CellML model to the CellML editorial board so that they can fill in the required ODE-based equations and metadata of this new CellML model.

### Load Discovered Models

As mentioned above, the user can make a shortlist of models from the discovered models for visualization, graphical editing, and model assembly and construction. Fig. 3 presents an example where the Biological meaning in Fig. 2 is replaced with Compartment, and Located in on the right of this example is added with a view to quickly provide more useful information to the user. In this space, we have provided some useful options: “view” option to get a detailed overview of a model; “delete” option to remove models if the user is not interested in considering these models on the platform; “visualization” option to find a similarity between a set of selected models; and “epithelial platform” option to navigate to the SVG-based interface and to semantically display the shortlisted models, which is discussed below.

**Figure 3.**
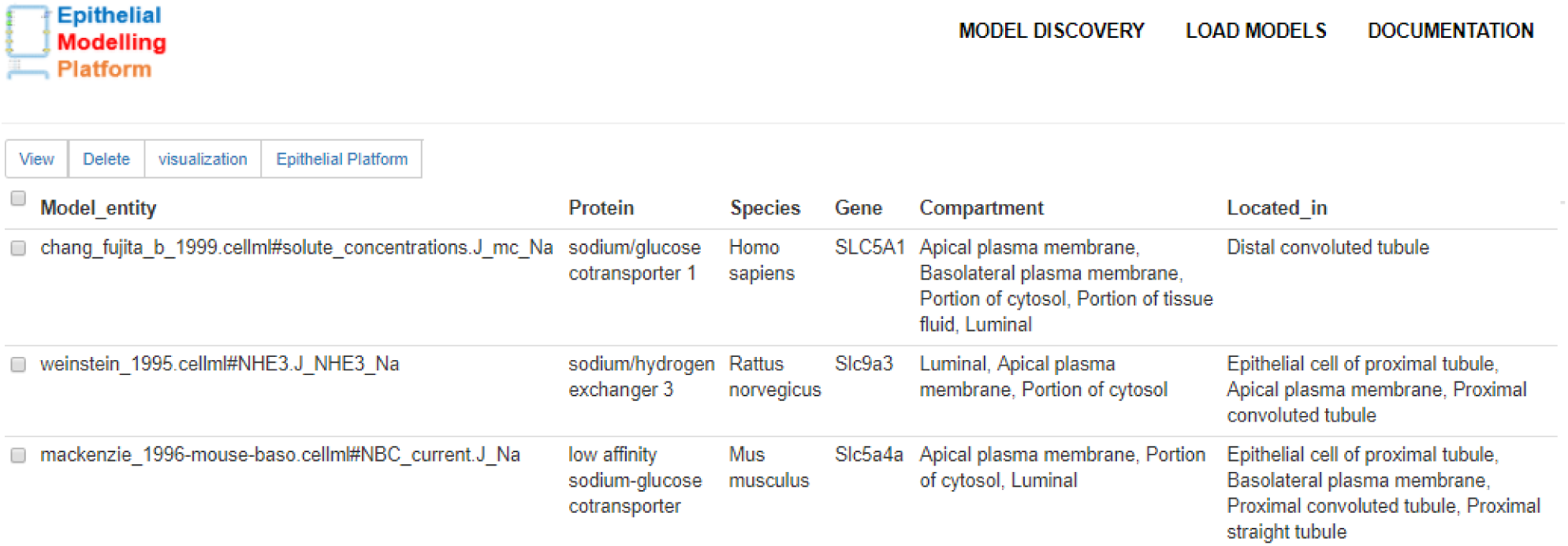
List of models for visualization and graphical editing on the platform. These models are selected from the model discovery interface illustrated in Fig. 2.

### Models of Similarity

Discovering similarity between models is vital to investigate coupling point(s), whereas visualization quickly conveys this investigation to the user. For this, we have developed models of similarity feature in the EMP. We aim to visually identify similar components, such as species, genes, and compartments between the selected models from the shortlist of models space. Fig. 4 shows a similar component between the NHE3 (Weinstein, 1995) and SGLT2 (Mackenzie et al., 1996) models. In this example, both models have same compartment. Nodes and edges are represented with unique colors to make the model distinguishable. Overall, by using this feature, the user would be able to analyze models and could choose relevant models in order to construct an epithelial model.

**Figure 4.**
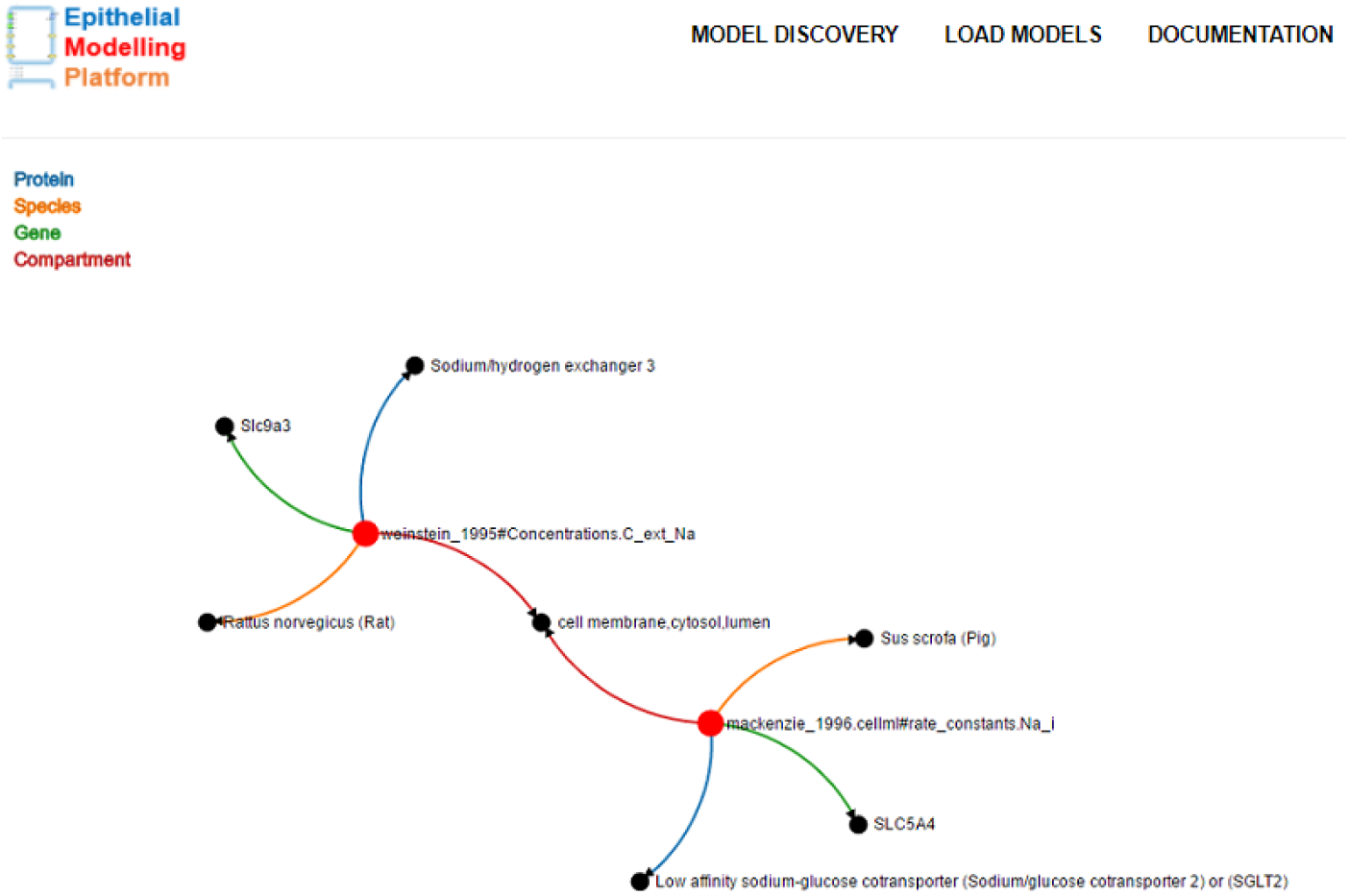
Illustrating the similarity between the NHE3 (Weinstein, 1995) and SGLT2 (Mackenzie et al., 1996) models. In this case, the transporter being modelled is located in the same compartment, labeled here with a unique colour. For convenience, we used “lumen” as a replacement of portion of renal filtrate in distal convoluted tubule and “cytosol” as a replacement of portion of cytosol in epithelial cell of distal tubule.

### Modelling Platform

Fig. 5 presents an example of our SVG-based model visualization, graphical editing, and model assembly and construction platform. This platform has five compartments: portion of renal filtrate in distal convoluted tubule (luminal compartment), portion of cytosol in epithelial cell of distal tubule (cytosol compartment), paracellular, portion of tissue fluid in epithelial cell of distal tubule (interstitial fluid compartment), and blood capillary; and three membranes: apical, basolateral, and capillary. These compartments and membranes have been annotated with unique colors as shown in the top-right corner of this example. In order to distinguish between fluxes and channels, we have used various shapes, as described in Figure 5.

The paracellular compartment separates two epithelial cells where diffusive fluxes are semantically placed on this compartment. In a similar manner, fluxes and channels are placed on a membrane based on the biological semantics. Before placing these fluxes and channels on a membrane, the platform checks its mediator (either apical, basolateral, or capillary membrane ID, and a protein ID) and shape (circles for fluxes or polygons for channels). A co-transporter, for example, Na-Cl co-transporter (TSC) in Fig. 5, is constructed if both (Na^+^and Cl^−^) fluxes have a mediator in common. On the right, checkboxes are generated for the visualized fluxes and channels in order to allow the user to move them across the membranes.

Direction of a flux is represented with a line, an arrow symbol, and a text to define the solute being transported. This direction is defined in the biological information deposited in PMR with source and sink parameters. Here, source and sink parameters can be interchangeably either portion of renal filtrate in distal convoluted tubule, portion of cytosol in epithelial cell of distal tubule, or portion of tissue fluid in epithelial cell of distal tubule compartment. For instance, Fig. 5 illustrates that flux of sodium (J mc Na) and flux of chloride (J mc Cl) flow from portion of renal filtrate in distal convoluted tubule to portion of cytosol in epithelial cell of distal tubule compartment across apical plasma membrane and TSC co-transporter. In contrast, concentration of annotated CellML variables float in the respective compartments. For example, C m K floats in the portion of renal filtrate in distal convoluted tubule compartment and C s Na in the portion of tissue fluid in epithelial cell of distal tubule compartment.

**Figure 5.**
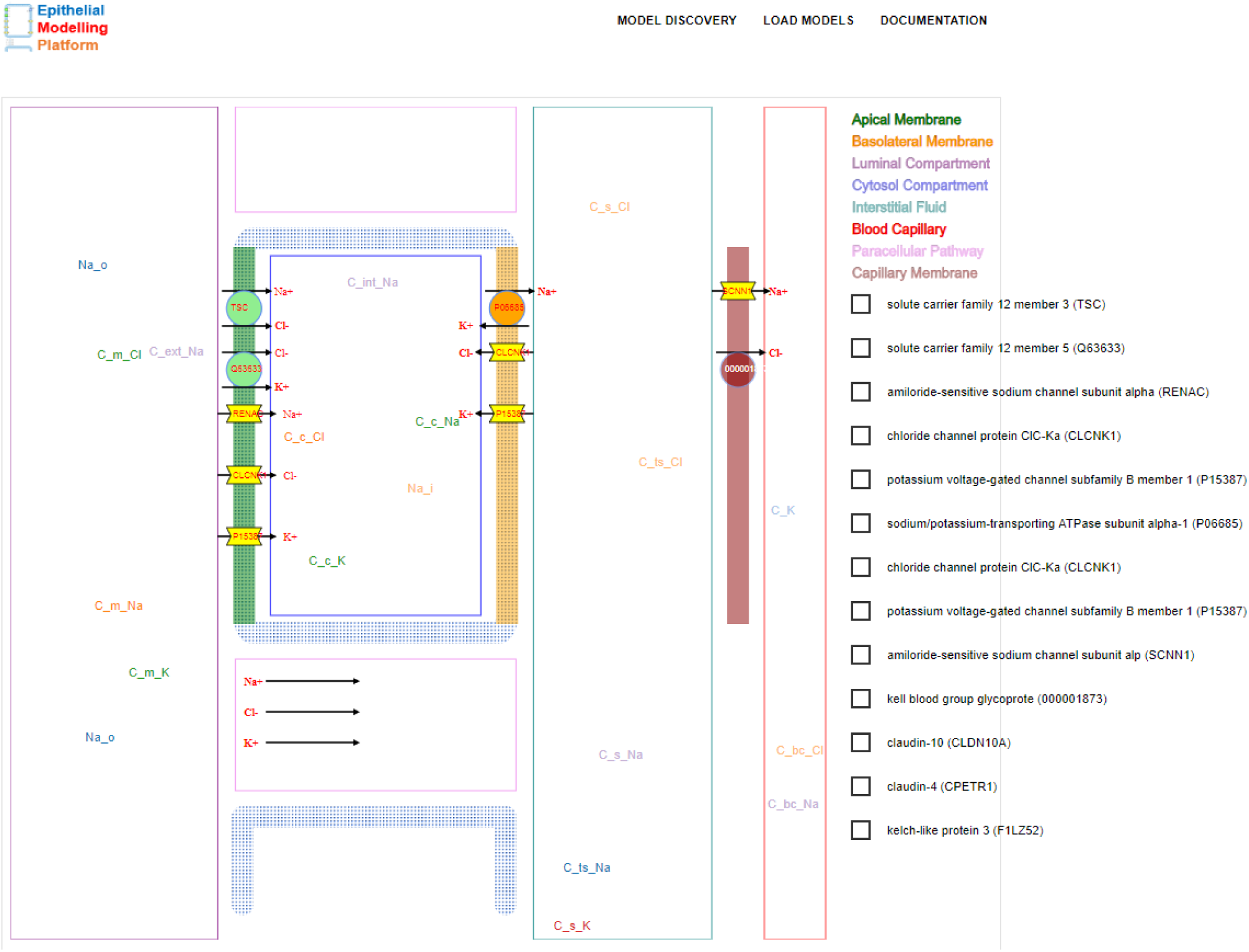
An example of the platform. It consists of five compartments and three membranes indicated by unique colors in the top-right corner. Physical entities and processes are represented with shapes such as: fluxes and co-transporters with circles; channels with polygons; and diffusive fluxes in the paracellular pathway with a text, a line, and an arrow. Arrows depict the direction of flow defined by the annotated biological semantics. On the right, checkboxes are generated for the visualized shapes (e.g. circles for fluxes, polygons for channels) on the apical, basolateral, and capillary membrane in order to allow the user to move across the membranes. Concentration of the annotated CellML variables float in the respective compartments. For example, C m K floats in the portion of renal filtrate in the distal convoluted tubule compartment and C s Na floats in the portion of tissue fluid in the epithelial cell of distal tubule compartment.

### Recommender System

On the SVG-based modelling platform of the EMP, we have provided a feature to make recommendations or suggestions for the user to perform graphical editing. In this case, users can drag a model from apical to basolateral membrane or vice versa, then a pop-up window will appear with a brief desription of the dragged model extracted from the biological semantics in PMR, as well as some suggestions provided by the EBI web services. Here, we have applied our concept presented in (Sarwar et al., 2018) in regard to find similar models as suggestions.

Similar to Figure 4 in (Sarwar et al., 2018), a list of suggestions begins with a brief description of the dragged model: CellML model entity with an URI pointing to the PMR workspace, species and gene used during the experiment, and a protein name. Next, this recommender system provides some suggestions on existing annotated models as opposed to the dragged model’s membrane. Fig. 6 shows an example when “flux of sodium from portion of renal filtrate in distal convoluted tubule to portion of cytosol in epithelial cell of distal tubule through the sodium/hydrogen exchanger 3 (NHE3) (Weinstein, 1995) model” is dragged from apical to basolateral membrane. Therefore, the recommender system extracts existing annotated models in the basolateral membrane from PMR. Users can choose one of the suggested models to replace with the dragged NHE3 model.

**Figure 6.**
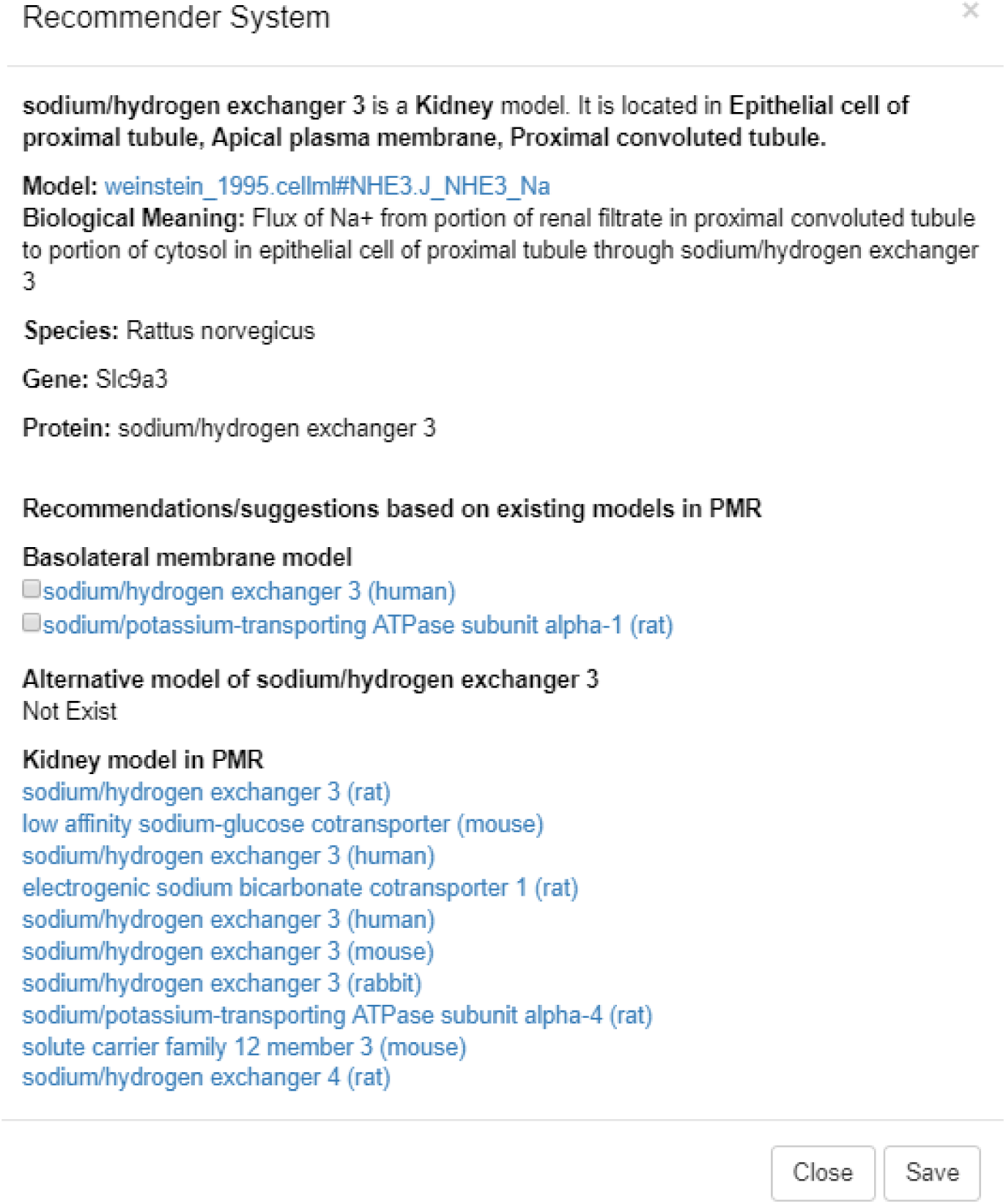
Recommender system as a pop-up window, with some information and suggestions, when a circle represented as “flux of Na^+^from portion of renal filtrate in distal convoluted tubule to portion of cytosol in epithelial cell of distal tubule through the sodium/hydrogen exchanger 3 (NHE3) (Weinstein, 1995) model” is moved from apical to basolateral membrane.

To populate the suggested models, we have leveraged two web services hosted at the EBI: WSDbfetch (WSDbfetch, 2019) and Clustal Omega (Clustal, 2019; Sievers et al., 2011). The WSDBfetch was used to get protein sequences for a list of protein identifiers. These protein sequences were formatted as JSON and then dispatched to the Clustal Omega web service to get a comparison score between the protein sequences in a representation of a similarity matrix. Fig. 7 illustrates a similarity matrix retrieved from the EBI Clustalo Omega web service.

**Figure 7.**
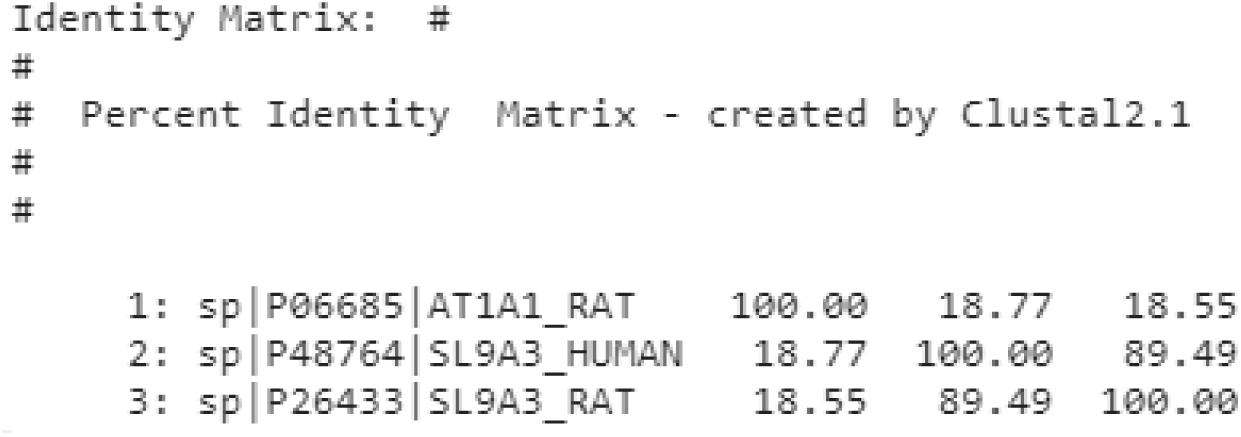
A similarity matrix retrieved from the EBI clustalo omega web service where P26433 is the dragged protein model and the remaining protein models are from Fig. 6, see list of proteins under “Basolateral membrane model” section header. Ranking of these proteins are based on IDs and matrix scores: P48764 (89.49) and P06685 (18.55), with respect to the dragged P26433 protein model (NHE3).

On the pop-up window, another recommendation is made as an alternative model of the dragged NHE3 model, i.e. same protein but different species, genes, or workspaces. In this case, “Not Exist” shows in Fig. 6. Lastly, a list of related organ models is shown for the users to investigate further.

### Model Assembly Service

The end result of our model discovery and assembly is a recipe that consists of models’ components and relevant information to help construct a new epithelial model. To achieve this, we have implemented a model assembly service which does model composition by utilizing the libCellML (libCellML, 2019) library. The source code of the model assembly service is available at https://github.com/dewancse/epithelial-modelling-platform/tree/master/server.

### libCellML

libCellML is a library to serialize, validate, and instantiate a CellML model by utilizing convenience methods and objects in C++. It also provides Python bindings to access these convenience methods and objects in Python. The source code and documentation of libCellML are available at https://github.com/cellml/libcellml and http://libcellml.readthedocs.io/.

#### Model-recipe and Composition

The first step of the model assembly service was to iterate over the model entities in the model-recipe with a view to extracting models’ components and then importing these components and the associated variables in the new model. Fig. 8 shows an example of eleven models’ components which is reproduced from Fig. 5. For example, Na-Cl co-transporter (TSC) is located on the apical membrane where “Na^+^and Cl^−^flow from portion of renal filtrate in distal convoluted tubule (“Luminal Compartment” indicated in the top-right corner in Fig. 8) to portion of cytosol in epithelial cell of distal tubule (“Cytosol Compartment” indicated in the top-right corner in Fig. 8) compartment”; and Na-K pump (P06685) is located on the basolateral membrane where “Na^+^flows from portion of cytosol in epithelial cell of distal tubule to portion of tissue fluid in epithelial cell of distal tubule (“Interstitial Fluid” indicated in the top-right corner in Fig. 8) compartment” and “K^+^flows from portion of tissue fluid in epithelial cell of distal tubule to portion of cytosol in epithelial cell of distal tubule compartment”. Relevant concentration of solutes float in the portion of renal filtrate in distal convoluted tubule, portion of cytosol in epithelial cell of distal tubule, and portion of tissue fluid in epithelial cell of distal tubule compartment.

**Figure 8.**
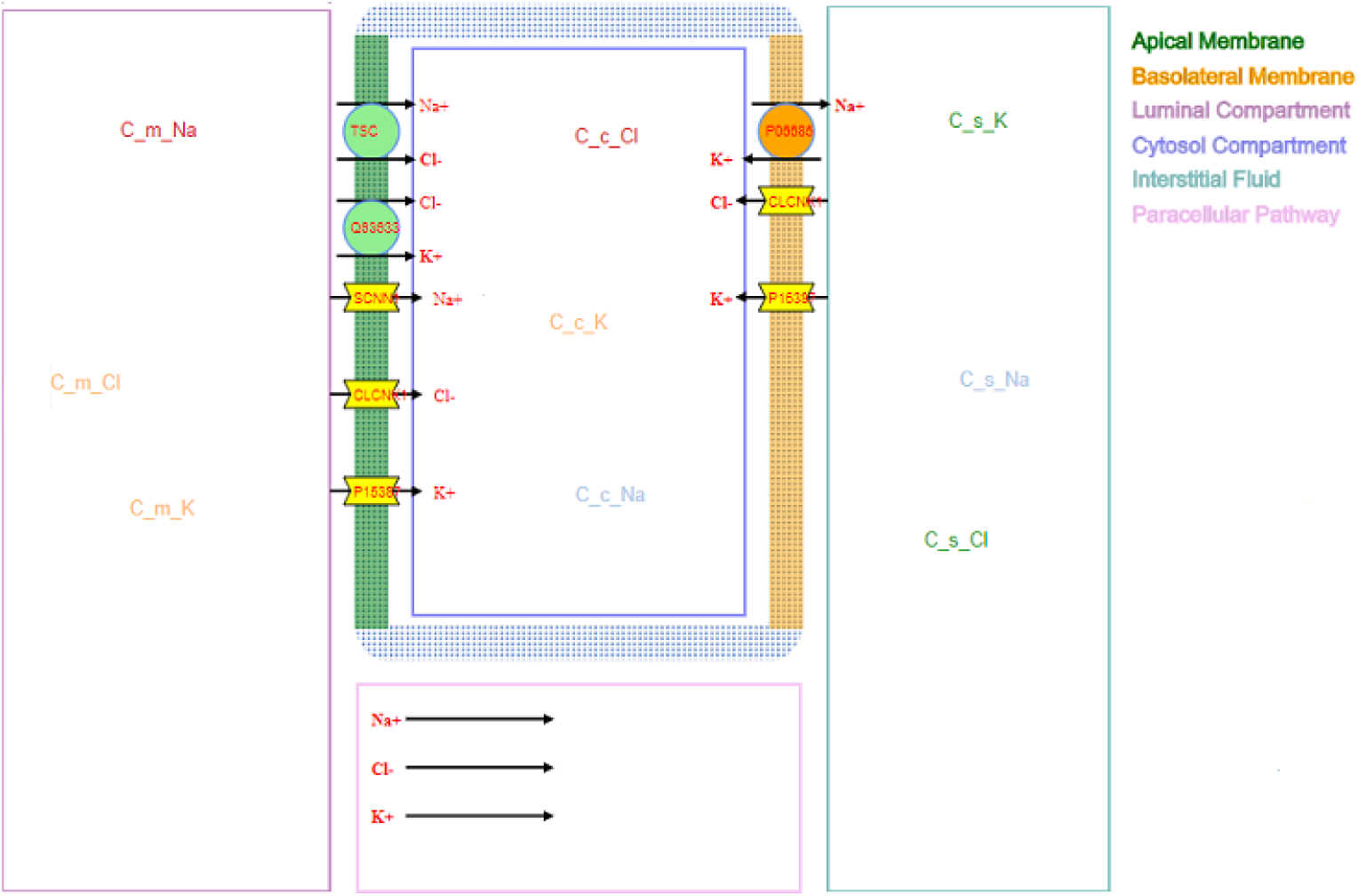
An example of the platform where Na-Cl and K-Cl co-transporters, Na channel, K channel, and Cl channel are located on the apical membrane; Na-K pump, K channel, and Cl channel are located on the basolateral membrane; and Na, K and Cl diffusive fluxes are located on the paracellular compartment. Respective concentration of solutes float in the portion of renal filtrate in distal convoluted tubule (“Luminal Compartment”), portion of cytosol in epithelial cell of distal tubule (“Cytosol Compartment”) and portion of tissue fluid in epithelial cell of distal tubule (“Interstitial Fluid”) compartment.

Fig. 9 shows a model-recipe, which is the background information of the visualized models in Fig. 8. This recipe has a detailed annotated information to construct a new epithelial model. In Fig. 9, med fma denotes mediator FMA which has the URI of apical plasma membrane; med pr denotes solute carrier family 12 member 3 or thiazide-sensitive sodium-chloride co-transporter; med pr text and med pr text syn denote the textual representation and synonym (in short form) of the med pr, respectively; model entity combines model name (chang fujita b 1999.cellml), component name (total transepithelial sodium flux) and variable name (J mc Na); model entity2 is the second part of this co-transporter where model name is chang fujita b 1999.cellml, component name is solute concentrations and variable name is J mc Cl; protein name, in this case, is a reference URI of the epithelial cell.

**Figure 9.**
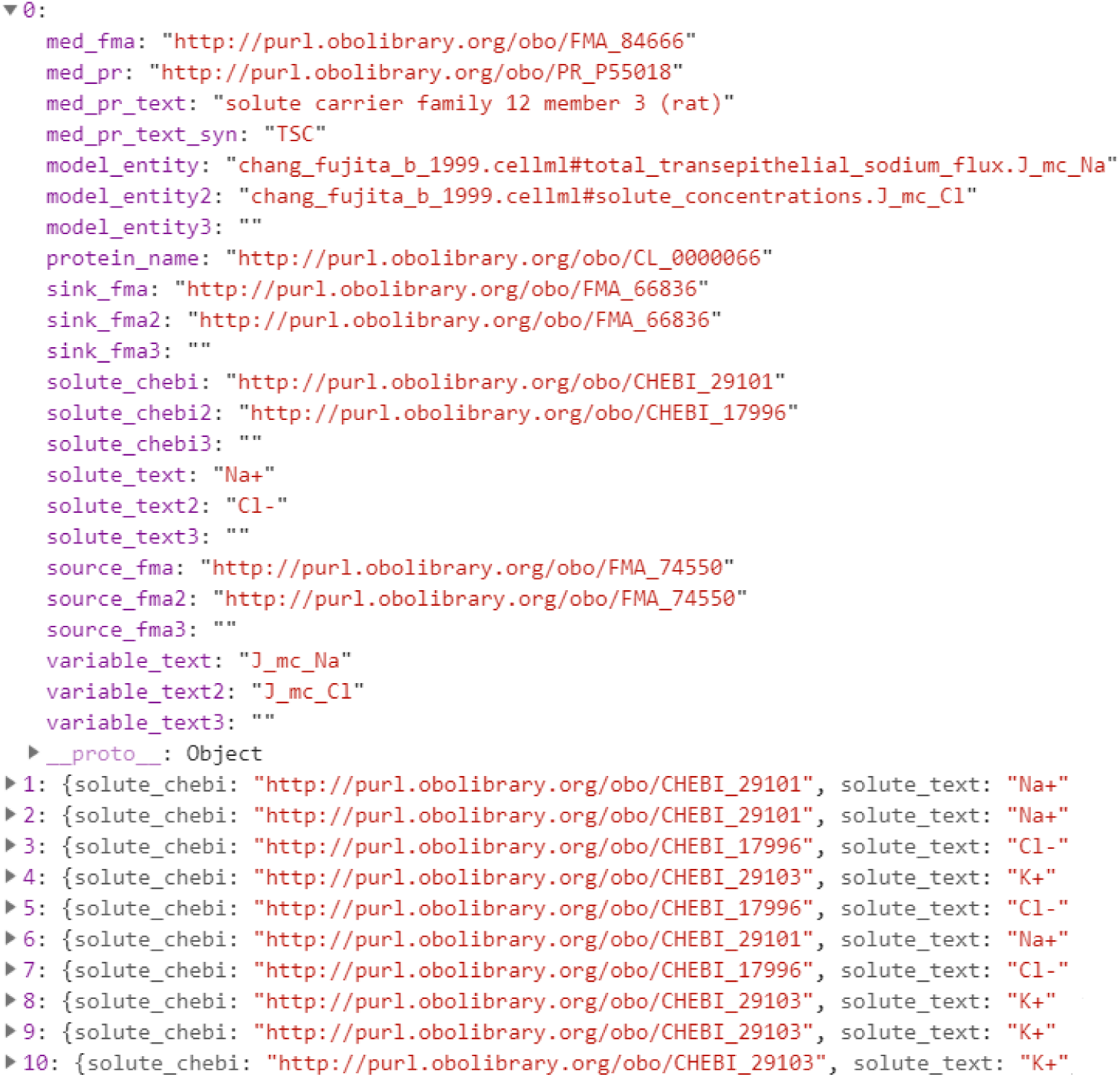
A detailed annotated information encapsulated inside a Na-Cl co-transporter (TSC) object visualized on the apical membrane in Fig. 8, which is a recipe to perform model composition by utilizing the libCellML (libCellML, 2019). The remaining co-transporters and channels visualized in Fig. 8 are kept hidden for convenience.

As shown in Fig. 8, “Na^+^and Cl^−^flow from portion of renal filtrate in distal convoluted tubule to portion of cytosol in epithelial cell of distal tubule compartment across apical plasma membrane and Na-Cl co-transporter (TSC)”, thus, source fma is a reference URI of portion of renal filtrate in distal convoluted tubule compartment, sink fma is a reference URI of portion of cytosol in epithelial cell of distal tubule compartment, solute chebi is a reference URI of sodium, solute text is a short form of sodium, variable text is a CellML variable name of sodium, source fma2 is a reference URI of portion of renal filtrate in distal convoluted tubule compartment, sink fma2 is a reference URI of portion of cytosol in epithelial cell of distal tubule compartment, solute chebi2 is a reference URI of chloride, solute text2 is a short form of chloride, and variable text2 is a CellML variable name of chloride. Similarly, Na-K pump in Fig. 8 has a detailed annotated information which is kept hidden in Fig. 9 for convenience.

In addition to importing components and the associated variables from the source models in the model-recipe, we encapsulated the imported components inside an epithelial component. Next, we created an environment component that contains a time variable, and also created three components - lumen (portion of renal filtrate in distal convoluted tubule), cytosol (portion of cytosol in epithelial cell of distal tubule), and interstitial fluid (portion of tissue fluid in epithelial cell of distal tubule) to store relevant variables from the model-recipe. The epithelial component contains additional variables that are required in the lumen, cytosol, and interstitial fluid component.

Following step involves iterating over the model entities in the model-recipe and then extracting variable and component names. As mentioned above, we encapsulated the extracted components inside an epithelial component. On the other hand, variable names were used to represent fluxes, channels, and diffusive fluxes. We constructed ordinary differential equations (ODE) to represent fluxes. However, for channels and diffusive fluxes, we extracted the full mathematical equations from the source models in the model-recipe. To construct these equations, we instantiated a time variable in the lumen, cytosol, and interstitial fluid component.

To construct an ODE equation, we need to have concentration of solutes in the respective compartments-portion of renal filtrate in distal convoluted tubule, portion of cytosol in epithelial cell of distal tubule, or portion of tissue fluid in epithelial cell of distal tubule. For example, to construct an ODE equation for “flux of sodium from portion of renal filtrate in distal convoluted tubule to portion of cytosol in epithelial cell of distal tubule compartment across apical plasma membrane”, we need to have “concentration of sodium in the portion of renal filtrate in distal convoluted tubule” and “concentration of sodium portion of cytosol in epithelial cell of distal tubule” compartment, then we can make an ODE equation which will measure “flux of sodium from portion of renal filtrate in distal convoluted tubule to portion of cytosol in epithelial cell of distal tubule” compartment, i.e. “sodium flows from portion of renal filtrate in distal convoluted tubule to portion of cytosol in epithelial cell of distal tubule” compartment. To get the respective concentration of solutes, we made a SPARQL call to PMR for a given solute (sodium, chloride, etc) and compartment (portion of renal filtrate in distal convoluted tubule, portion of cytosol in epithelial cell of distal tubule, or portion of tissue fluid in epithelial cell of distal tubule). The result of this call was a list of concentration of the specified solute in the specified compartment. Then we iterated over each concentration variable in the result and checked that whether both the flux and concentration variable reside in the same component. In the ODE equation, we marked with plus or minus sign notation to denote in which direction the concentration of the solute flows. After getting the respective concentration variable of the flux and the notation, we constructed an ODE equation by utilizing a MathML (MathML, 2019). Fig. 10 illustrates an ODE equation -concentration of sodium in the portion of renal filtrate in distal convoluted tubule compartment (C m Na) with respect to time is equal to negative flow of concentration of sodium (J mc Na). This flow is negative because concentration of sodium is moving from portion of renal filtrate in distal convoluted tubule to portion of cytosol in epithelial cell of distal tubule compartment.

**Figure 10.**
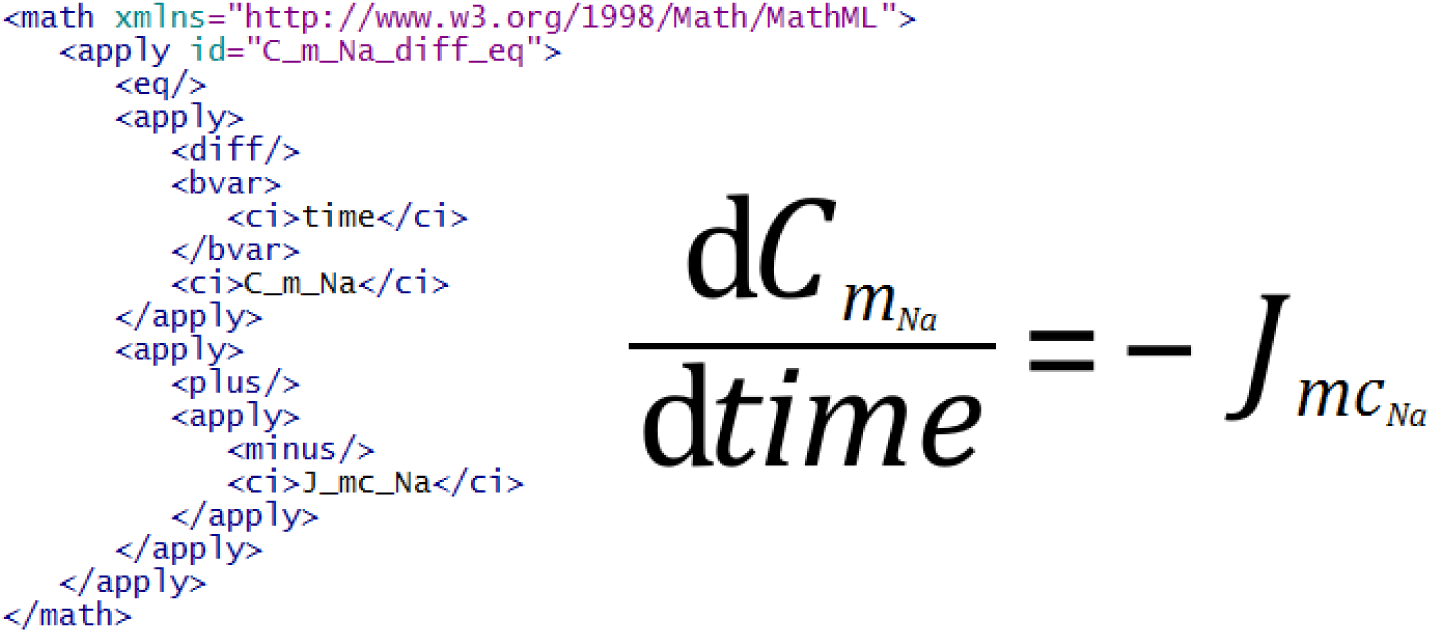
A MathML (MathML, 2019) representation of “flux of sodium” (J mc Na) which is an ordinary differential equation for “concentration of sodium” (C m Na) with respect to time.

In contrast, to construct mathematical equations for channels and diffusive fluxes, we extracted the full math equation from the source model in the model-recipe. For example, Fig. 11 shows an example equation of a sodium channel from the Chang and Fujita model (Chang and Fujita, 1999). We extracted this channel’s equation and traversed through the variables of this equation in order to instantiate them in the epithelial component of the new model.

**Figure 11.**
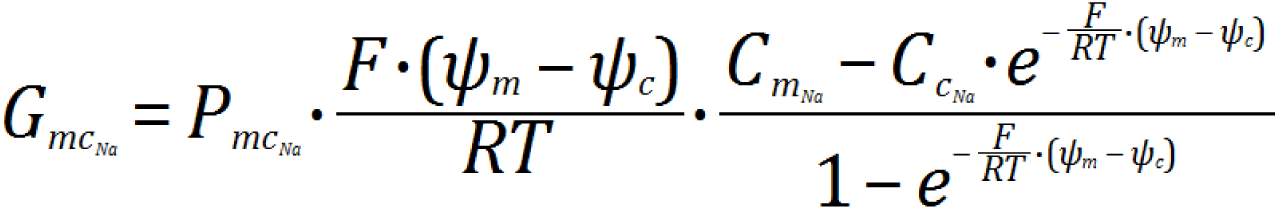
Equation of a “sodium channel” (G mc Na) visualized as polygons (SCNN1) on the apical membrane in Fig. 8.

In terms of total fluxes, channels and diffusive fluxes equations, we evaluated out-going and in-coming fluxes of components and then constructed equations. For example, Fig. 12 illustrates “total flux of sodium” and “total flux of chloride” in the portion of renal filtrate in distal convoluted tubule (luminal compartment) compartment of Fig. 8. In the final step of the model assembly service, we mapped variables between the epithelial component and its encapsulated components. We also mapped variables between epithelial component’s encapsulated siblings.

**Figure 12.**
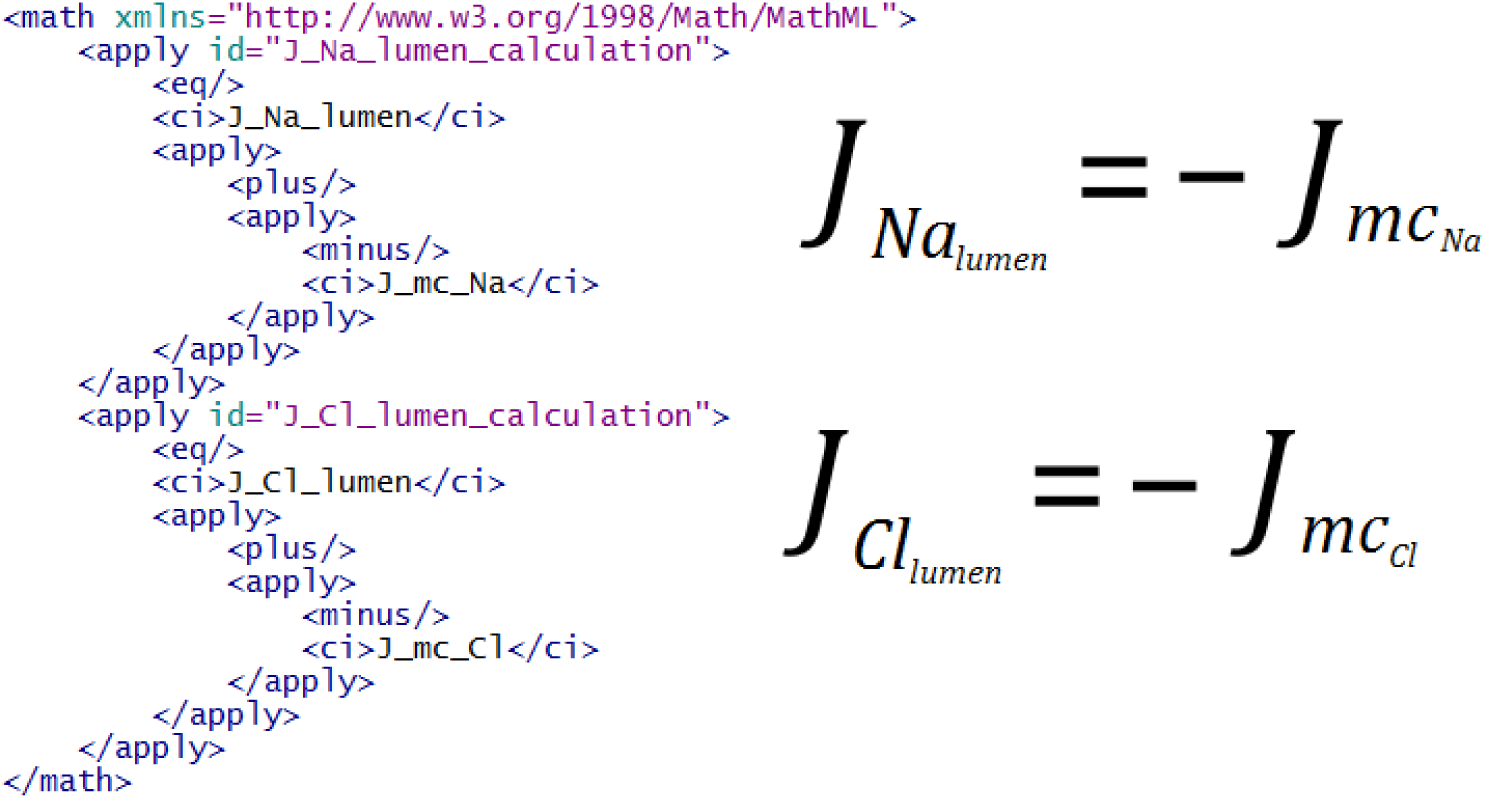
A MathML (MathML, 2019) representation of “total flux of sodium” (J Na lumen) and “total flux of chloride” (J Cl lumen) in the portion of renal filtrate in distal convoluted tubule (Luminal Compartment) compartment in Fig. 8.

## 4 DISCUSSION

We have developed a novel web-based platform, the Epithelial Modelling Platform (EMP), for scientists to discover and explore mathematical models relevant to their work. On this platform, they can semantically visualize models for graphical editing and model assembly with a view to constructing a new model. In particular, this involves three steps: annotation of mathematical models with biological information; generation of metadata using the SemGen tool; and the EMP to discover models from the metadata in order to visualize models for graphical editing and model assembly and construction.

To perform biological annotation, we have comprehensively annotated a cohort of epithelial transport models: twelve renal and four lung models, which are available at https://models.physiomeproject.org/workspace/527. In particular, we pulled out mathematical terms from each model and annotated with biological information. In addition to this, we curated the models with protein identifiers from the UniProt database, species and gene names used during the experiments of the models, compartment names, and anatomical locations. Next, we fed these biological information onto the SemGen tool to generate metadata. This metadata is then deposited in PMR which are discoverable, accessible and persistent.

To discover and explore the epithelial transport models, we have developed the EMP that utilizes cascaded SPARQL calls to iteratively retrieve information from PMR. First, we make a query with a protein identifier to the OLS in order to get a human-readable name. Subsequent call retrieves protein’s species and gene name. In addition to model discovery, users are able to add protein models and generate metadata in PMR. The EMP allows users to semantically display a collection of discovered models for graphical editing and model assembly. This “graphical editing” has been integrated with a recommender system where users can rank similar models based on the suggestions provided by the EBI web services.

The end result of the graphical editing and model assembly is the generation of a model-recipe, which is a collection of components from disparate discovered models. Fig. 9 illustrates an example model-recipe that has the required information to perform model composition. This recipe is then dispatched to our model assembly service in order to compose a new epithelial model. To perform model composition, we have utilized convenience methods and objects of the libCellML library.

The integration and discovery of the semantic annotation visualized on the SVG-based epithelial platform is a novel application of the services provided by PMR that would inspire users to reuse the data and models. Integration of data and models with the provision of semantic web technologies will integrate computational model of various organ systems, which is the ultimate effort of the Physiome Project and the VPH. We believe this approach with the aid of modern tools and technologies would provide an excellent epithelial modelling platform. Novice modellers could use this platform as a learning tool.

The “Add Model” feature enables scientists to create a new CellML model and metadata. In the future, we would like to extend this feature and implement a notification system between CellML editorial board and the scientists so that the CellML editorial board can incorporate mathematical equations and validate the new CellML model.

## FUNDING

DMS was supported by the Medical Technologies Centre of Research Excellence’s Doctoral Scholarship. DPN was supported by an Aotearoa Foundation Fellowship. JHG, BEC, and MLN were supported by the National Institutes of Health grant R01LM011969.

### ACKNOWLEDGMENTS

The authors wish to acknowledge the Centre for eResearch at the University of Auckland for their help in facilitating this research. The availability of our live web application demonstration is made possible by use of the Nectar Research Cloud, a collaborative Australian research platform supported by the National Collaborative Research Infrastructure Strategy (NCRIS). Thanks to Sean Matheny at the Center for eResearch for his cooperation to setup our tool and service in Center for eResearch at the University of Auckland and Nectar.

## AVAILABILITY OF DATA AND MATERIALS

The source code and links to the live demonstration of the Epithelial Modelling Platform are available at https://github.com/dewancse/epithelial-modelling-platform.

